# Guano morphology reveals ecological information in British bats

**DOI:** 10.1101/728824

**Authors:** Roselyn Lydia Ware, Benjamin Garrod, Hannah Macdonald, Robin G Allaby

## Abstract

Bats are primary consumers of nocturnal insects, disperse nutrients across landscapes, and are excellent bioindicators of an ecosystem’s health, however four of the seventeen Great British species are listed as declining. In this study we aim to investigate the link between bat guano morphology and diet, specifically looking at the ability to predict 1) species, 2) dietary guild and 3) bat size, using guano morphology alone. It was found that guano morphology overlapped too much to make predictions on species however, in some cases, it could be used to predict dietary guild or size.

## 2. Introduction

Bats (order Chiroptera) are the second largest group of mammals. Bats are pivotal to supporting global biodiversity; they are the primary consumers of nocturnal insects, disperse nutrients across landscapes, and are excellent bioindicators of an ecosystem’s health (Jones et al., 2009, Patterson et al., 2003). However, 25% of bats worldwide are classed as ‘of conservation concern’, with a further 21% classified as ‘near threatened’ (Boyles and Storm, 2007). Four of the seventeen Great British species are listed as declining, with the status of several others unknown (I.U.C.N., 2019). The plethora of threats faced by bats include (but are not limited to): unsympathetic development projects, destruction of tree lines and hedgerows, the drainage of wetlands, infectious diseases, and the impact of pesticides (Ashrafi et al., 2011, Mickleburgh et al., 2002). Additionally, climate change may have a highly detrimental impact on bats, including changes in prey abundances, alterations in the efficacy of echolocation calls, and the consequences of extreme weather events (Luo et al., 2014). This is why it is vital to understand their ecological niches and correctly identify species.

Using bat guano to detect an individual’s diet using either the stable isotope method or the molecular approach has been well established in its accuracy and application (Clare et al., 2009, Hope et al., 2014, Salvarina et al., 2013). However, it does involve a large budget and in-depth knowledge of the subject. In this study, we wished to investigate whether any potentially useful ecological information, such as diet, could be determined by studying guano morphology alone. The hypothesis being that different dominant prey species may influence the size and shape of the guano produced. To test this hypothesis, we compiled diets from all 17 bat British bat species (*B. barbastellus, E. serotinus, M. bechsteinii, M. brandtii, M. daubentonii, M. mystacinus, M. nattereri, N. lesleri, N. noctule, P. auratus, P. austriacus, P. nathusii, P. pipistrellus, P. pygmaeus, R. ferrumequinum, R. hipposideros* and M. *alcathoe*) and compared this with a data set of guano morphology. In 1997, Vaughan undertook a review of all of the published diets of the 15 species of bats then known to be present in Great Britain (Vaughan, 1997). Since then, two new species of bat have been described, and have been identified to be present in Great Britain: *P. pygmaeus* and *M. alcathoe*. Furthermore, numerous additional studies have since been published. It was, therefore, deemed valuable to produce an up-to-date synthesis of the all data pertaining to the diets of Great British bats.

In this study we investigate guano morphology data’s ability to identify 1) species, 2) dietary guilds and 3) bat size. Information on species, dietary guild, or size may be used to give ecologists some useful information in the field about bat ecology. Understanding dietary overlaps, and the basis of the diets, will be valuable in helping to direct conservation efforts, and in explaining inter-species interactions (Patterson et al., 2003, Schnitzler et al., 2003).

## 3.0. Materials and methods

### 3.1. Diet

The diet data used in this study was collected from a comprehensive search of the available literature, searching exclusively for bats present in Britain (*B. barbastellus, E. serotinus, M. bechsteinii, M. brandtii, M. daubentonii, M. mystacinus, M. nattereri, N. lesleri, N. noctule, P. auritus, P. austriacus, P. nathusii, P. pipistrellus, P. pygmaeus, R. ferrumequinum, R. hipposideros*, and *M. alcathoe*). Google Scholar and the Web of Knowledge databases were mined for published bat diets up to January 2016. The terms searched for using the Boolean “AND” were: “Diet”, “Food”, and “Prey”, with “Bat”, “Chiroptera”, and the names of each of the species. In some cases, it was appropriate to exclude certain studies. Typically, this was due to the reporting of diets with only presence/absence data, as it was not possible to convert these to numerical data without the introduction of bias. Papers were also excluded if they did not detail diet break down; if they described controlled experiments (i.e. captive fed bats) or they did not use primary data.

In total, we identified 80 published studies, this resulted in a dataset of 215 diets spread across 17 species. For more information regarding the references, including country of sample collection, please see supplementary information, Table S1. For the purpose of this study, the data are expressed in number of diets rather than number of studies, as each study may present more than one diet. They may present the diets of different species; or present the diets of one species measured in different ways; or diets of one species from different sampling locations or seasons. When directly comparing with guano morphology *M. alcathoe* was dropped from the dataset as there was no guano samples available with which to compare.

### 3.2. Guano Morphology

Guano samples were collected, and to ensure that the samples were as representative of each species as possible, samples were selected to cover the whole of Great Britain (as far as the range of the species allowed). In addition to Stebbings’ diagnostic characteristics of length (minimum – maximum within a sample), diameter (minimum to maximum) and particle size (Stebbings, 1986); we measured colour, presence/absence of nodulation, presence/absence of tip points, and presence/absence of curvature. The criterion for categorising particle size and colour can be found in the supporting information, Table S2. Nodulation, tip pointing and curvature were observed by eye and recorded as presence/absence data.

To ensure correct species identification, after measurement guano samples were identified by DNA barcoding. Individual guano samples were crushed and incubated overnight at 37°C in 300 μl CTAB buffer on a sample agitator at 400rpm. CTAB extractions yield higher concentrations of DNA from guano than MoBio, Epicentre, and Qiagen stool kits (Jedlicka et al., 2013). After incubation DNA was extracted using chloroform:isoamyl alcohol 24:1. After spinning, the DNA is in the aqueous phase, and proteins and polysaccharides move into the chloroform/alcohol layer, removing these inhibitors. The DNA was then purified using DNeasy columns and buffers, with an additional acetone wash and dry before elution (Köchl et al., 2005, Prado et al., 1997).

The species of bat was confirmed using barcoding as follows; 20 μl PCRs were prepared using a mixture of all of the primers shown in Table S3, supplementary material, each at 5 μM. Each PCR contained 2 μl 10X Platinum^®^ Taq buffer, 2 μl of dNTPs at 2mM, 0.8μl 50mM Mg++, 1.3 μl primer mix, 0.1 μl Platinum^®^ Taq DNA polymerase, between 0.2-2 μl of sample and 11.8-13.6 μl ultrapure H2O. Touchdown PCR was used in order to account for the differences in optimum annealing temperatures of the primers used (Don et al., 1991, Korbie and Mattick, 2008). Touchdown thermal cycling conditions were as follows: 5 mins at 95°C, followed by 10 cycles of 94°C for 30s, 57°C for 30s (decreasing by 0.1°C per cycle) and 72°C for 30s, followed by 32 cycles of 95°C for 30s, 54°C for 30s, then 72°C for 30s, followed by a final extension period of 72°C for 7 minutes.

After PCR, success was determined by running on a 2% agarose gel, stained with gel red. Clean-up was undertaken by adding 2 μl of Fast-AP and 0.5 μl of Exonuclease-1, then incubated at 37°C for 30mins, then 80°C for 15 mins. Forward primers (BF1-7) were used in a GATC LightrunTM Sanger sequencing reaction. Sequences were checked from traces using CodonCode aligner, then sequences were confirmed using the NCBI database.

This process resulted in a sample size of 104 positivity identified samples from the 16 target species to compare with the diet dataset. A summary of sample sizes per species for both the diet and guano datasets is available in supplementary information, Table S4.

### 4.0. Statistical analysis

In order to compare between diet and guano datasets, Principle Component Analysis (PCA) was undertaken using R function prcomp (Holland, 2008) as this gave the highest proportion of variance in the first components. PCA plots comparing principle component 1 (PC1) and principle component 2 (PC2) were created using R and data was assigned by 1) species, 2) dietary guild and 3) size of bat. To assign dietary guilds diets were sorted using complete-linkage clustering using the hclust package in R, dependant on diet composition. To assign size categories the minimum and maximum weight per bat species were gathered from the Bat Conservation Trust (http://www.bats.org.uk/pages/ukbats.html#Resident) and plotted to visualise size groupings (please see supplementary information, Figure S1).

Spatial autocorrelation within PCA plots was calculated using the Moran.I function in the ape v5.3 R package (Gittleman and Kot, 1990) to assess whether the clustering of the assigned groups was significant.

Wilcoxon sign rank test was conducted comparing PC1 for diet to PC1 for guano, PC2 v PC2 etc. This allowed us to use the individual sample dataset retaining the variation within each grouping and look at one proposed cluster at a time. This method was used to compare clusters assigned by 1) species, 2) dietary guild and 3) size.

### 5.0. Results

Figures 1a and 1b show PCA plots showing clusters assigned to species for diet (Figure 1a) and guano (Figure 1b). Principle component 1 represents 58.24% and 54.83% of the Proportion of Variance for diet and guano, respectively, with PC1 and PC2 cumulatively represent 83.39% and 70.46% of the data for diet and guano, respectively. For diet, the Moran’s I analysis when the data is assigned to species shows significant clustering (p = 2.510526e-05), this was expected because the diet data carries the assumption that the same species will have a similar diet, therefore this result is a product of the method. Clustering the guano data by species also yielded a significant result (p = 3.352874e-14). This was investigated further with Wilcoxen signed rank test comparison of PC1, which represents the highest proportion of variance for both datasets (58.24% and 54.83% for diet and guano data respectively). This analysis revealed 9 species out of the 16 had a significant correlation between diet and guano morphology in PC1 (*B. barbastellus, E. serotinus, M. daubentonii, M. nattereri, P. auritus, P. austriacus, P. nathusii, P.pipistrellus*, and *P.pygmaeus*) (see Table S5a in supplementary information for full results).

**Figure 1.a-f.**
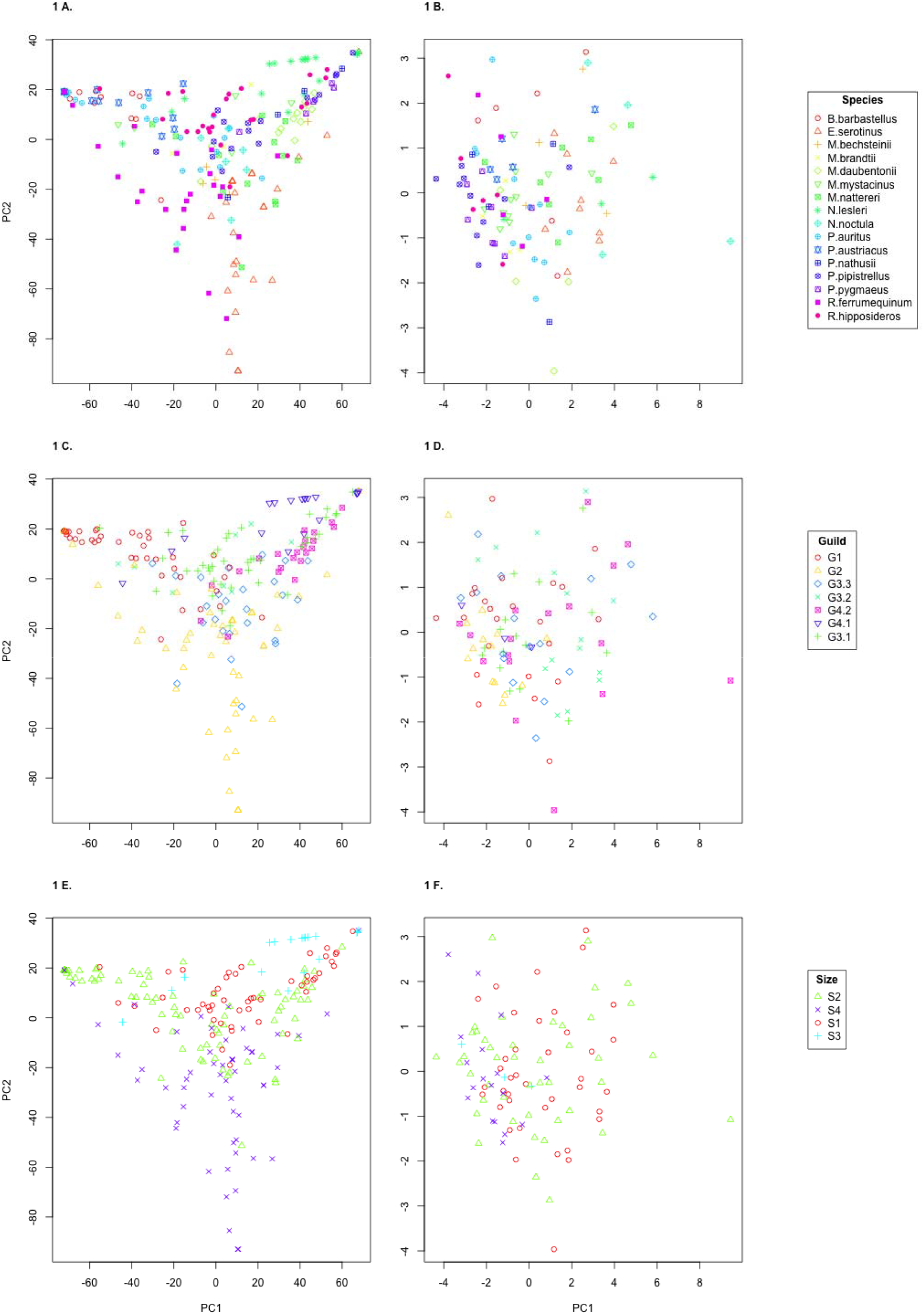
Principle Component Analysis plots of both Diet and Guano data assigned by Species (A,B), Guild (C,D) and Size (E,F).

The data was then grouped into dietary guilds (Figure 2) with guild 1 (G1) composed of bat species with a diet of largely Lepidoptera; Guild 2, bat species with diets containing a largely Coleoptera; Guild 3 from found from complete-linkage clustering was broken down further into G3.1, generalist and Diptera diet; G3.2, generalist, Lepidoptera and Diptera diet and G3.3, generalist with Coleoptera. Guild 4 from complete-linkage clustering was broken down to G4.1, a diet consisting of almost all Diptera, and G4.2 a diet consisting of largely Diptera with notable Trichoptera presence.

**Figure 2.**
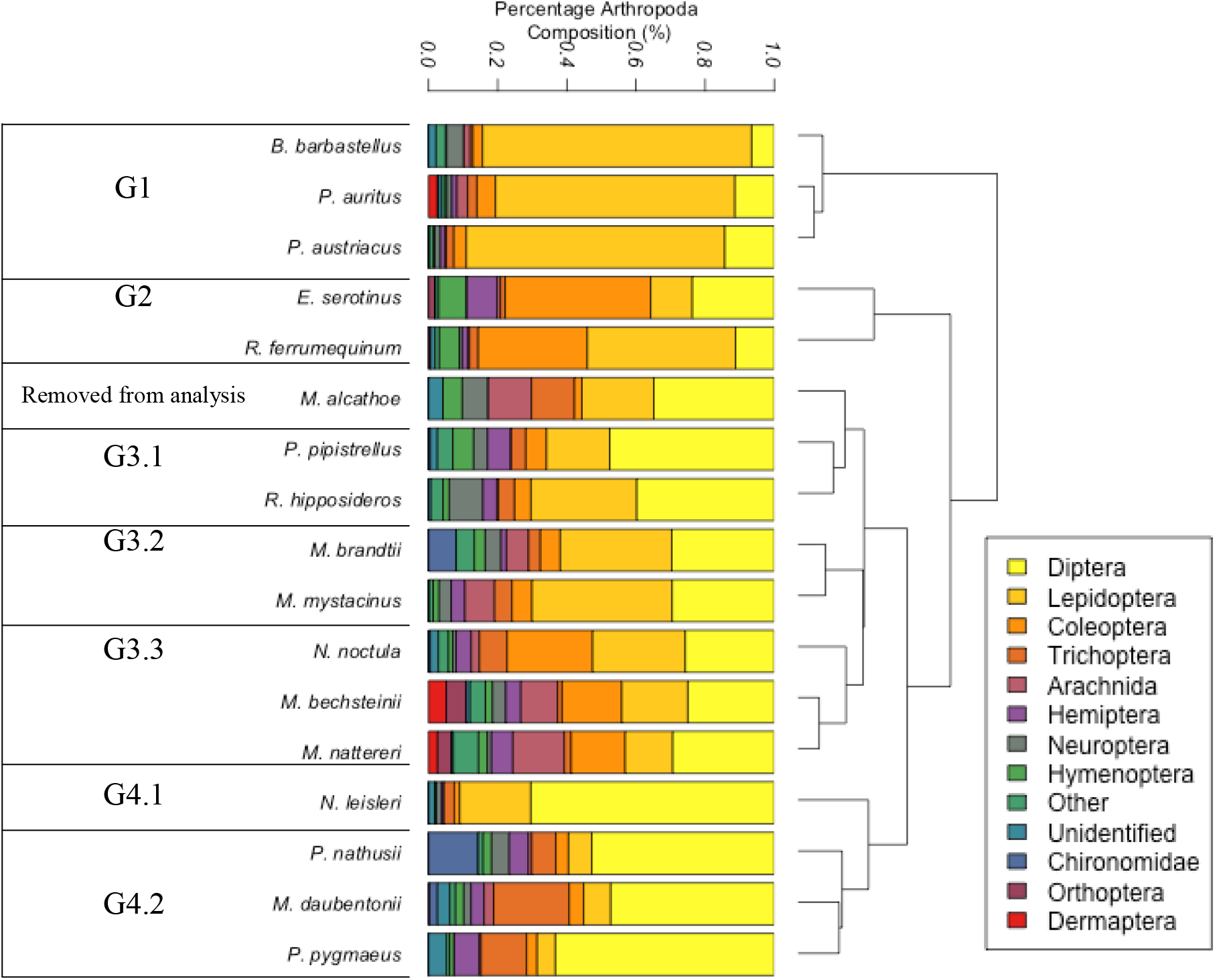
The diet of each bat species from the literature. The average diets of each of the species using all of the methods. Prey taxa have been grouped by order (where possible), class, phylum, or kingdom (where necessary). Diets sorted using complete-linkage clustering using hclust in R.

Figure 1c and d, show PCA plots assigned to dietary guilds (defined in Figure 2) for diet (Figure 1c) and guano (Figure 1d). The clustering of guilds within the diet data is highly significant (p = <0.001) which is exactly as expected because the guilds were assigned using the diet data. However, interestingly the guilds also produce significant clustering within the guano data (p = 0.0156828). Wilcoxen’s signed rank test comparison revealed this was mainly driven by G1, G3.1, and G4.2 (please see Table S5b in supplementary for full results).

Figures 1e and 1f show PCA plots assigned to bat size (defined in Figure S1 Supplementary Information) for diet (Figure 1e) and guano (Figure 1f). Moran’s I analysis shows the clustering for both diet and guano was highly significant (p = <0.001) when assigned by size, somewhat matching the distributions within dietary guild. Wilcoxen’s signed rank test comparison revealed all four size groupings were significantly correlating when comparing PC2vPC2 but only Size category S1, the smallest bat species were significant when comparing PC1 of both data sets (please see Table S5b in Supplementary Information for full results).

## 6.0. Discussion

The aim of this study was to identify whether bat guano morphology data could be used to predict bat diet. We first compared bat diet and bat guano morphology among the all species. As expected, the guano morphology overlapped too substantially to be able to separate all 16 species of bat compared. It is nevertheless interesting that nine species when investigated individually do seem to correlate between diet and guano morphology. Figure 2b shows it is possible to distinguish a few species from each other for example, *P. pipistrellus* rarely overlaps with *E. serotinus*. Generally, however, it would be impossible to distinguish between for example *M. daubentonii, P. austriacus, P. nathusii*, and *P. auritus* as they all overlap considerably in terms of guano morphology.

However, grouping the data by dietary guilds shows significant clusters within the guild morphology data. Though the clusters still overlap there is a pattern within the data particularly between Guild 1 (diet consisting of largely Lepidoptera), Guild 3.1, (generalist and Diptera diet), Guild 3.3 (generalist with Coleoptera) and Guild 4.2, (largely Diptera with notable Trichoptera presence), where species following these diets generally produce similar size and shape of guano. When looking at grouping by the size of the bat, there is a similar pattern to that shown in dietary guild, with the largest separation between the smallest (S1) species and the largest (S4) species. Considering that the size and diet of a bat is intrinsically linked with one another (Barclay et al., 1991) it would be hard to pull apart their separate influences on the guano morphology data. A species within the smallest group of bats is more likely to have a very different diet than a species in the largest group of bats, for example, *P. pipistrellus* and *R. hipposideros* are both in S1 and G3.1 and are quite different to *E. serotinus* and *R. ferrumequinum* who are both in S4 and G2, therefore it is unknown the true cause of the divergence.

When taking into consideration the density (i.e. where the majority of the samples fall) and the limitations of the study, this data could be used to predict the dietary guild and size of a bat. This may have applications within outreach or preliminary ecological assessment for field ecologists and be used to potentially identify certain characteristics of the environment, for example, the morphology of a guano sample could be used to tell the researcher whether the sample came from a Coleoptera rich environment, or what invertebrate species may be important to the particular bat in question.

Predicting the dietary guild of a species present in an area is interesting and informative because typically, the most important factors in bat niche separation are considered to be the partitioning of habitat and diet (Ashrafi et al., 2011, Schoener, 1974). The ranges of many of the Great British bat species are acutely overlapping (Bat Conservation Trust, 2009), suggesting that trophic resource partitioning is important in supporting the species in Great Britain (Aguirre et al., 2002, Arlettaz et al., 2000). Prey availability may, in some circumstances, be a more important determiner of species ability to co-inhabit in an area than dietary differences (Griffiths, 1975). If a prey species is rare, it may become a limiting factor, thus introducing competition between those predating upon it. If a prey species is common, competitive exclusion can cause one bat species to be excluded (Hardin, 1960).

In general, bats with a higher extinction risk are those with greater dietary specialisation (Boyles and Storm, 2007, Jones et al., 2003). Where a bat has a narrow dietary range, it is considered more vulnerable to extinction, whereas a bat with a broad dietary breadth is considered to be more robust. Knowledge of the level of dietary specialisation of a species, coupled with intelligence of the conservation status of the prey species, is a valuable resource.

