## Supplemental Tables and Figures for "Guano morphology reveals ecological information in British bats"

### Supplementary Information

***Table S1.*** *References used to characterise diets, including species studied and country of sample collection.*

| Reference | Year of publication | Species studied | Country of sample collection | Reference |
|---|---|---|---|---|
| Ahmim & Moali | 2013 | *R. ferrumequinum, R. hipposideros* | Algeria | (Ahmim and Moali, 2013) |
| Andreas | 2010 | *P. auritus, P. austriacus* | Czech Republic | (Andreas, 2010) |
| Andreas et al. | 2013 | *R. ferrumequinum, R. hipposideros* | Slovakia | (Andreas et al., 2013) |
| Andreas et al. | 2012 | *B. barbastellus* | Czech Republic | (Andreas et al., 2012b) |
| Andreas et al. | 2012 | *M. bechsteinii, M. nattereri, P. auritus* | Central Europe | (Andreas et al., 2012a) |
| Arlettaz et al. | 2000 | *P. pipistrellus, R. hipposideros* | Switzerland | (Arlettaz et al., 2000) |
| Ashrafi et al. | 2011 | *P. auritus, P. austriacus* | Switzerland | (Ashrafi et al., 2011) |
| Barlow | 1997 | *P. pipistrellus, P. pygmaeus* | Britain | (Barlow, 1997) |
| Bárta | 1975 | *P. auritus* | Slovak Republic | (Bárta, 1975) |
| Bartonička et al. | 2008 | *P. pygmaeus* | Czech Republic | (Bartonička et al., 2008) |
| Bauerová | 1982 | *P. austriacus* | Czech Republic | (Bauerova, 1982) |
| Bauerová & Cerveny | 1986 | *M. nattereri* | Czech Republic | (Bauerová and Cerveny, 1986) |
| Beck | 1995 | *B. barbastellus, E. serotinus, M. mystacinus, N. leisleri, N. noctula, P. auritus, P. austriacus, P. nathusii, P. pipistrellus, R. ferrumequinum* | Switzerland | (Beck, 1995) |
| Beck | 1994 | *R. ferrumequinum* | Switzerland | (Beck et al., 1994) |
| Beck | 1991 | *M. nattereri* | Switzerland | (Beck, 1991) |
| Beck et al. | 1989 | *R. hipposideros* | Switzerland | (Beck et al., 1989) |

| | | | | |
|---|---|---|---|---|
| Bontadina et al. | 2008 | *R. hipposideros* | Switzerland | (Bontadina et al., 2008) |
| Boonman | 1995 | *P. auritus, R. ferrumequinum* | Netherlands, Belgium | (Boonman, 1995) |
| Buckhurst | 1930 | *P. auritus* | Britain | (Buckhurst, 1930) |
| Catto et al. | 1994 | *E. serotinus* | England | (Catto et al., 1994) |
| Chung et al. | 2015 | *E. serotinus* | Korea | (Chung et al., 2015) |
| Danko et al. | 2010 | *M. alcathoe* | Slovakia | (Danko et al., 2010) |
| Duvergé | 1997 | *R. ferrumequinum* | Britain | (Vaughan, 1997) |
| Feldman et al. | 2000 | *P. austriacus, R. hipposideros* | Israel | (Feldman et al., 2000) |
| Flanders & Jones | 2009 | *R. ferrumequinum* | Britain | (Flanders and Jones, 2009) |
| Flavin et al. | 2001 | *M. daubentonii* | Ireland | (Flavin et al., 2001) |
| Gajdosik & Gaisler | 2004 | *E. serotinus* | Czech Republic | (Gajdosik and Gaisler, 2004) |
| Gerber et al. | 1994 | *E. serotinus* | Switzerland | (Gerber et al., 1994) |
| Gloor et al. | 1989 | *N. noctula* | Switzerland | (Gloor et al., 1989) |
| Hanson | 1950 | *P. auritus* | Sweden | (Vaughan, 1997) |
| Heinicke & Krauβ | 1978 | *P. auritus* | Germany | (Heinicke and Krauß, 1978) |
| Hoare | 1991 | *P. pipistrellus* | England | (Hoare, 1991) |
| Hollyfield | 1993 | *M. mystacinus, P. auritus, R. hipposideros* | Ireland | (Vaughan, 1997) |
| Hope et al. | 2014 | *M. nattereri* | England | (Hope et al., 2014) |
| Jin et al. | 2005 | *R. ferrumequinum* | China | (Jin et al., 2005) |
| Jing et al. | 2010 | *R. ferrumequinum* | China | (Wang Jing, 2010) |
| Jones | 1995 | *N. noctula* | Britain | (Jones, 1995) |
| Jones | 1990 | *R. ferrumequinum* | China | (Jones, 1990) |
| Kaňuch et al. | 2005 | *N. noctula, N. leisleri* | Slovakia, Czech Republic | (Kaňuch et al., 2005a) |
| Kaňuch et al. | 2005 | *N. leisleri* | Slovakia | (Kaňuch et al., 2005b) |

| | | | | |
|---|---|---|---|---|
| Kervyn & Libois | 2008 | *E. serotinus* | Belgium | (Kervyn and Libois, 2008) |
| Krauss | 1978 | *P. auritus* | Germany | (Krauss, 1978) |
| Kruger et al. | 2013 | *M. daubentonii* | Germany | (Krüger et al., 2013a) |
| Kruger et al. | 2013 | *P. nathusii* | Latvia | (Krüger et al., 2013b) |
| Kruger et al. | 2012 | *M. daubentonii* | Germany | (Krüger et al., 2012) |
| Leishman | 1983 | *R. ferrumequinum, R. hipposideros* | Britain | (Vaughan, 1997) |
| Lino et al. | 2014 | *R. hipposideros* | Portugal | (Lino et al., 2014) |
| Lucan et al. | 2009 | *M. alcathoe* | Czech Republic | (Lucan et al., 2009) |
| Ma et al. | 2008 | *R. ferrumequinum* | China | (Ma et al., 2008) |
| Mackenzie & Oxford | 1995 | *N. noctula* | Britain | (Mackenzie and Oxford, 1995) |
| Manwaring-Banes | 1939 | *P. auritus* | Britain | (Manwaring, 1939) |
| McAney & Fairley | 1989 | *R. hipposideros* | Ireland | (McAney and Fairley, 1989) |
| McAney et al. | 1991 | *E. serotinus* | Czech Republic | (McAney, 1991) |
| Mikula & Čmoková | 2012 | *E. serotinus* | Czech Republic | (Mikula and Čmoková, 2012) |
| Nissen et al. | 2013 | *M. daubentonii* | Germany | (Nissen et al., 2013) |
| Oldfield | 1990 | *P. auritus* | Sweden | (Vaughan, 1997) |
| Pir | 1994 | *R. ferrumequinum* | Luxembourg | (Vaughan, 1997) |
| Poulton | 1929 | *R. ferrumequinum* | Britain | (Poulton, 1929) |
| Ransome | 1996 | *R. ferrumequinum* | Britain | (Vaughan, 1997) |
| Razgour et al. | 2011 | *P. auritus, P. austriacus* | Britain | (Razgour et al., 2011) |
| Robertson | 1988 | *R. ferrumequinum* | Britain | (Vaughan, 1997) |
| Robinson | 1990 | *P. auritus* | Britain | (Robinson, 1990) |

| | | | | |
|---|---|---|---|---|
| Robinson & Stebbings | 1993 | *E. serotinus* | Britain | (Robinson and Stebbings, 1993) |
| Rostovskaya et al. | 2000 | *P. auritus* | Central Russia | (Rostovskaya et al., 2000) |
| Roswag et al. | 2015 | *M. bechsteinii, M. nattereri, P. auritus* | Central Germany | (Roswag et al., 2015) |
| Rydell | 1989 | *P. auritus* | Sweden | (Rydell, 1989) |
| Rydell et al. | 1996 | *B. barbastellus* | Germany, Sweden | (Rydell et al., 1996) |
| Siemers & Swift | 2006 | *M. bechsteinii, M. nattereri* | Germany | (Siemers and Swift, 2006) |
| Shiel et al. | 1991 | *M. nattereri, P. auritus* | Ireland | (Shiel et al., 1991) |
| Shiel et al. | 1998 | *N. leisleri* | Ireland, England | (Shiel et al., 1998) |
| Sierro & Arlettaz | 1997 | *B. barbastellus* | Switzerland, Asia | (Sierro and Arlettaz, 1997) |
| Smirnov & Vekhnik | 2014 | *E. serotinus, M. brandtii, M. daubentonii, M. mystacinus, M. nattereri, N. leisleri, N. noctula, P. auritus, P. nathusii, P. pipistrellus, M. daubentonii* | Russia | (Smirnov and Vekhnik, 2014) |
| Sologor | 1980 | *E. serotinus* | Ukraine | (Sologor, 1980) |
| Sullivan et al. | 1993 | *M. daubentonii, N. leisleri, P. pipistrellus* | Ireland | (Sullivan et al., 1993) |
| Swift & Racey | 1983 | *M. daubentonii, P. auritus* | Scotland | (Swift and Racey, 1983) |
| Swift et al. | 1985 | *P. pipistrellus* | Scotland | (Swift et al., 1985) |
| Taake | 1993 | *M. bechsteinii, M. brandtii, M. daubentonii, M. mystacinus, M. nattereri, P. auritus* | Germany | (Taake, 1993) |
| Thompson | 1982 | *P. auritus* | England | (Thompson, 1982) |
| Vesterinen et al. | 2013 | *M. daubentonii* | Finland | (Vesterinen et al., 2013) |

| | | | | |
|---|---|---|---|---|
| Walhovd & Hoegh- Gildberg | 1984 | *P. auritus* | Denmark | (Vaughan, 1997) |
| Waters et al. | 1999 | *N. leisleri* | Britain | (Waters et al., 1999) |
| Waters et al. | 1995 | *N. leisleri* | Britain | (Waters et al., 1995) |
| Whitaker | 1994 | *M. nattereri, P. austriacus* | Israel | (Vaughan, 1997) |
| Whitaker & Karataş | 2009 | *B. barbastellus, E. serotinus, M. brandtii, M. mystacinus, M. nattereri, P. auritus, P. austriacus, P. pipistrellus, P. pygmaeus, R. ferrumequinum, R. hipposideros* | Turkey | (Whitaker Jr and Karatas, 2009) |
| Williams et al. | 2011 | *R. hipposideros* | Britain | (Williams et al., 2010) |
| Wolz | 1993 | *M. bechsteinii* | Germany | (Vaughan, 1997) |
| Zeale et al. | 2011 | *B. barbastellus, M. nattereri, P. pipistrellus* | Britain | (Zeale et al., 2011) |
| Zukal & Gajdošík | 2012 | *E. serotinus* | Czech Republic | (Zukal and Gajdošík, 2012) |

***Table S2.*** *Measuring colour (a) and particle size (b)*

a) Particle size

| | |
|---|---|
| 1. | Very fine, smooth outline, small divots |
| 2. | Fine, mostly smooth outline, bigger divots |
| 3. | Medium size, rough outline, medium divots |
| 4. | Quite coarse, rough outline, medium to large divots |
| 5. | Coarse, very rough outline, large divots |

b) Colour

| | |
|---|---|
| 1. | Light to dark yellow |
| 2. | Light brown with yellow flecks present |
| 3. | Light-medium brown |
| 4. | Medium-dark brown with some black flecks |
| 5. | Over half of the guano is black |

***Table S3.*** *Primers used to confirm the identity bat species of the guano*

a) Forward

| Primer name | Orientation | Sequence |
|---|---|---|
| BF1 | Forward | ATGACAAACAYTCGAAAATCC |
| BF2 | Forward | ATGACAAACATTCGAAAGTMC |
| BF3 | Forward | ATGACCAACATTCGTAAATCW |
| BF4 | Forward | ATGACCAACATTCGAAAATCY |
| BF5 | Forward | ATGACCMACATTCGAAAATCY |
| BF6 | Forward | ATGACCAACATTCGAAAGTCY |
| BF7 | Forward | ATGACCAACATTCGCAARTCY |
| BX1 | Reverse | GTCTGMTGTRTAGTGTATGG |

b) Reverse

| | | |
|---|---|---|
| BX2 | Reverse | RTCYGATGTGTGATGCATGG |
| BX3 | Reverse | RTCTGATGTRTAGTGTATTGC |
| BX4 | Reverse | RTCTGATGTRTARTGTATGGC |
| BX5 | Reverse | RTCTGAYGTRTAGTGTATAGC |
| BX6 | Reverse | RTCTGATRTGTAATGTATAGC |
| BX7 | Reverse | ATCTGATGTATAATGTATWGCT |
| BX8 | Reverse | GTCTGATGTATAGTGTATGGA |
| BX9 | Reverse | GTCTGGTGTGTAATGTATGG |
| BX10 | Reverse | ATCTGATGTAGTGCGCATGG |

***Table S4.*** *Table summarising sample sizes for a) each species, b) each dietary guild and c) each size category.*

a) Species

| **Species** | **Guano sample size** | **Diet sample size** |
|---|---|---|
| *Barbastellus barbastellus* | 6 | 11 |
| *Eptesicus serotinus* | 9 | 26 |
| *Myotis bechsteinii* | 4 | 5 |
| *Myotis brandtii* | 4 | 3 |
| *Myotis daubentonii* | 5 | 13 |
| *Myotis mystacinus* | 11 | 5 |
| *Myotis nattereri* | 8 | 13 |
| *Nyctalus leisleri* | 3 | 19 |
| *Nyctalus noctula* | 4 | 7 |
| *Plecotus auritus* | 12 | 26 |
| *Plecotus austriacus* | 5 | 11 |
| *Pipistrellus nathusii* | 4 | 4 |
| *Pipistrellus pipistrellus* | 10 | 16 |
| *Pipistrellus pygmaeus* | 8 | 5 |
| *Rhinolophus ferrumequinum* | 5 | 23 |
| *Rhinolophus hipposideros* | 6 | 24 |
| *Myotis alcathoe* | - | 4 |
| **Total** | **104** | **215** (211 without *M. alcathoe*) |

b) Guild

| Guild | Diet | Guano |
|---|---|---|
| G1 | 48 | 23 |
| G2 | 49 | 14 |
| G3.1 | 41 | 16 |
| G3.2 | 8 | 15 |
| G3.3 | 25 | 16 |
| G4.1 | 19 | 3 |
| G4.2 | 22 | 17 |

c) Size

| Size class | Diet | Guano |
|---|---|---|
| S1 | 51 | 39 |
| S2 | 86 | 44 |
| S3 | 19 | 3 |
| S4 | 56 | 18 |

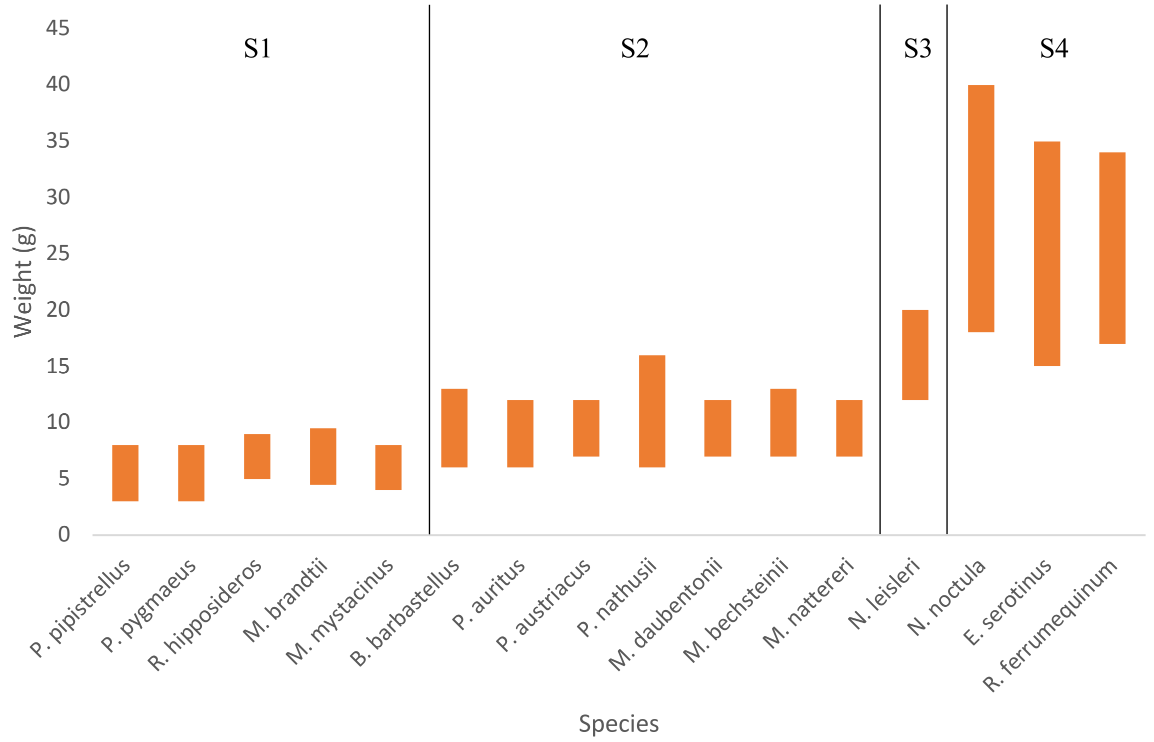


**Figure S1.** Assigning size categories to British bats using minimum and maximum weight from. Weights gathered per species from the Bat conservation trust website (http://www.bats.org.uk/pages/uk_bats.html#Resident)

***Table S5.*** *Showing the results of Wilcoxen signed rank test comparing the PCA outputs of Diet and Guano morphology. Only* significant correlations are presented:
* p-value = <0.05, ** p-value = <0.01, *** p-value = <0.001.

a) Species

| Species | Component 1 | Component 2 | Component 3 | Component 4 |
|---|---|---|---|---|

| | | | | |
|---|---|---|---|---|
| Diet Data: Proportion of Variance | 58.24% | 25.15% | 5.608% | 2.299% |
| Guano Data: Proportion of Variance | 53.83% | 15.63% | 13.59% | 11.56% |
| *B. barbastellus* | *** | *** | | |
| *E. serotinus* | *** | *** | | |
| *M. bechsteinii* | | | | |
| *M. brandtii* | | | | |
| *M. daubentonii* | * | | *** | ** |
| *M. mystacinus* | | | | |
| *M. nattereri* | * | | | ** |
| *N. lesleri* | | ** | | |
| *N. noctula* | | | | |
| *P. auritus* | ** | ** | | |
| *P. austriacus* | ** | ** | * | |
| *P. nathusii* | * | | | |
| *P. pipistrellus* | *** | | | |
| *P. pygmaeus* | ** | ** | | |
| *R. ferrumequinum* | | ** | | |
| *a* | | ** | | |

b) Guild

| **Guild** | **comp1** | **comp2** | **comp3** | **comp4** |
|---|---|---|---|---|
| G1 | *** | *** | * | |
| G2 | | *** | * | |
| G3.1 | ** | *** | | |
| G3.2 | | | | |
| G3.3 | | * | | * |
| G4.1 | | ** | | |
| G4.2 | *** | *** | *** | ** |

c) Size

| **Size** | **comp1** | **comp2** | **comp3** | **comp4** |
|---|---|---|---|---|
| S1 | *** | *** | | |
| S2 | | *** | | |
| S3 | | ** | | |
| S4 | | *** | ** | |

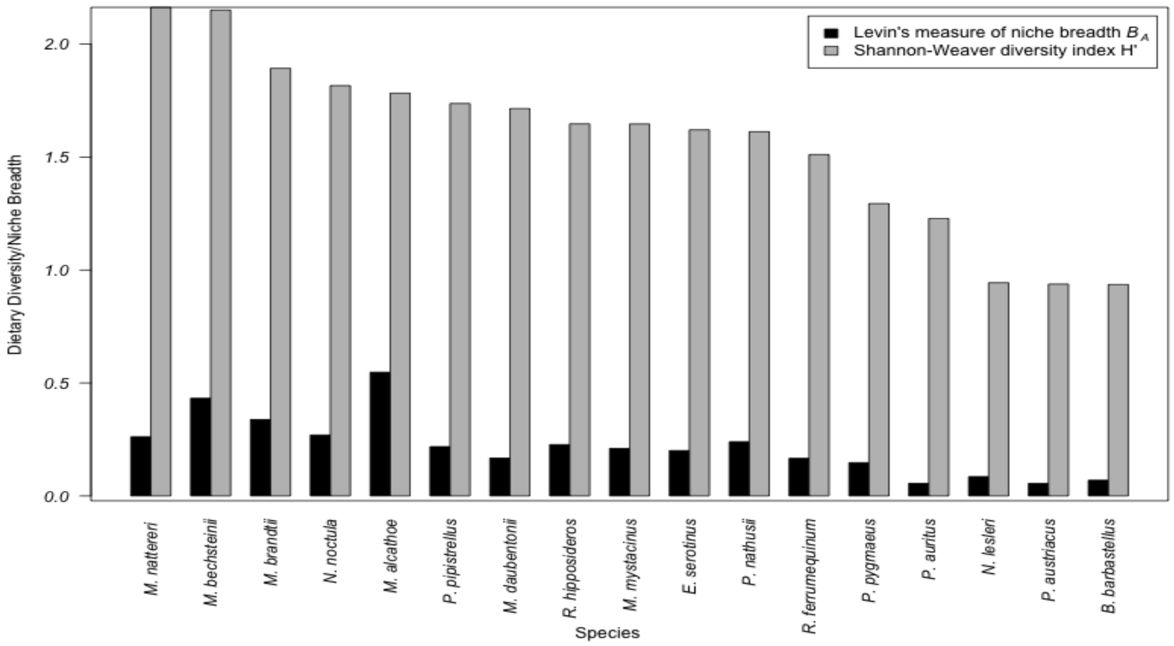


***Figure. S2.*** *Dietary diversity and niche breadth of each species.* The dietary diversity calculated using Shannon-Weaver diversity index (H') (grey) and niche breadth calculated using Levin's standardised index ($B_A$) (black).
